# Estimation of discovery rate in multiple-comparison tests at low sampling numbers

**DOI:** 10.1101/2025.02.23.639778

**Authors:** Marian Manciu

**Affiliations:** Physics Department, University of Texas at El Paso

**Keywords:** Bonferroni Correction, Multivariate Analysis, Multiple Comparions Procedures, Statistical Inference, Likelihood

## Abstract

The traditional statistical methods are very powerful if the sampling number in an experiment meets at least the minimum power analysis requirement. However, at lower sampling numbers, they tend to dramatically underestimate the discoveries leading to a large percentage of type II errors. Sometimes, the minimum sampling number is not reached, particularly in preliminary data in large-scale, simultaneously multiple-comparison experiments, where such a required number is much larger than that needed for one-comparison experiments. This work presents a simple approach for searching for discoveries in multiple-comparison experiments with low sampling numbers, based on the fact that the obtained *p*-values should be uniformly random when no discovery is present. While the proposed approach is less powerful than traditional methods at large sampling numbers, it maintains its efficiency at low sampling numbers. Thus, it is important for preliminary data analysis, for which traditional methods might not exhibit any statistically significant discoveries due to the low available sampling numbers. This approach helps to design further experiments with sufficient power for the desired significance level.

## 1. Introduction

High-throughput analysis, such as microarrays that measure expression levels of thousands of genes simultaneously, or high-throughput screening, which allows the performance of millions of chemical or genetic tests automatically, have become ubiquitous tools in many scientific disciplines. The large number of hypotheses tested simultaneously raises the possibility of large numbers of false discoveries (type I errors) to arise solely by chance. A family of statistical tests for multiple comparisons has been developed and applied to avoid these errors, such as Tukey’s honest significance difference criterion, Fisher’s least significant difference procedure, Dunn and Sidak’s approaches, Scheffe’s S procedure, and the most known, the Bonferroni correction [1].

The Bonferroni correction states that the null hypothesis should be rejected at the level α for one test only if *p< α/m*, where *m* is the number of simultaneous tests. This correction is considered too stringent for practical purposes of not-so-high sampling numbers. As a result, it has been relaxed by a sequential rejection procedure that was pioneered by Holm [2] and further improved by Hochberg [3], and by Benjamini and Hochberg [4]. Benjamini and Hochberg’s procedure remains the “golden standard” for very large numbers *m* of simultaneous tests (e.g., *m* of the order of 10^4^ for differential gene expressions), by controlling the False Discovery Rate (FDR) at a desired level *α (FDR < α)*.

The conventional approach of statistics is very powerful when it follows the framework for which it was designed. For example, for any desired level of significance (*α* being the acceptable percentage of false positive) and any desired detection power 1-*β* (*β* being the acceptable percentage of false negative), there is a minimum required number of samplings for which effects of desired magnitudes (i.e., discoveries) can be detected. This is also valid for the Bonferroni correction for (simultaneously) multiple comparison experiments, with the caveat that a larger number of samplings is required to achieve the desired precision, than in single comparison experiments.

Sampling is time-consuming, costly, and/or sometimes not available. The situation is worse when a large number *m* of comparisons, are tested simultaneously. For example, to apply successfully Bonferroni correction, the level of significance should be decreased to 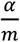, with *α* being the level of significance desired for a single comparison. If *m* = 10000, 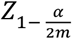 is two to four times larger than 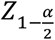,resulting in a needed increase in the sampling number by a factor of the order of 10 as compared to that of a single comparison experiment.

The focus of this paper is to suggest a simple method for estimating the number of discoveries at much lower sampling numbers than those required for applying the Bonferroni method. The accounting for an unusually high amount of low-p values is done typically via a family of methods known as *p-value histograms*, which were pioneered by Pyke [5]. Other well-known procedures suggested recently are the *q*-value method of Storey [6], the Beta-Uniform Mixture (BUM) model of Pounds and Morris [7], and the Spatial Local Regression Histogram (SPLOSH) of Pounds and Cheng [8]. Compared to these procedures, here a simpler method for detecting existing discoveries is suggested.

Our method considers that the distribution of the *p*-values obtained from the null hypotheses should be uniformly random. If such distribution is not uniformly random at the required level of significance, at least one null hypothesis needs to be removed and the process repeated. Although at large sampling numbers, our approach is less powerful than the standard methods of Bonferroni [1] and Benjamini and Hochberg [4], it has the advantage that its results depend much less on the sampling number. If discoveries exist, their cumulative signature will show up at low sampling numbers, as a deviation from the (expected) uniformly random distribution of the *p*-values.

The reason to look for an alternative approach for looking for discoveries in experiments performed at much lower sampling numbers than those required by traditional power analysis will be discussed first in the context of examining the results of a simple “thought” experiment. It will be shown that for such an experiment, whereas the traditional approaches cannot reject any null hypothesis, the overall likelihood of such a result (when all null hypotheses hold), is vanishingly small.

Next, we will examine the likelihood of the overall result, for the example provided in the seminal paper of Benjamini and Hochberg [4]. We will argue that, instead of looking for individual discoveries (e.g., null hypotheses, which can be rejected at the significance level desired), one might alternatively look for how likely is the result, if no “discoveries” would be present. In other words, the traditional statistics approach is to test the likelihood of individual hypotheses to not be null, and at low sampling numbers, almost none of the “null” hypothesis can be rejected. We suggest to look instead at the likelihood of the result if all null hypotheses would be true; and if the likelihood of the result is extremely small, at least one null hypothesis should not be valid.

In large-scale multiple comparison experiments, if all null hypotheses would hold, the *p* values obtained should be uniformly randomly distributed. Large divergences from that distribution are likely due to some null hypothesis not being true. From the wealth of algorithms employed to detect the departure from uniform random distribution, which were developed for verifying the quality of random number generators [9], we will restrict here to the oldest and best-known one, the Kolmogorov-Smirnov test for the Cumulative Distribution Function (CDF) [10].

Finally, the suggested approach for estimating the number of discoveries in multiple comparison experiments, will be compared with the Bonferroni and Benjamini-Hochberg methods for Monte-Carlo simulations of large-scale experiments. It will be shown that our approach has a good power to detect discoveries even at very low sampling numbers, much lower than those required for the Bonferroni and even Benjamini-Hochberg procedures, to be efficient.

## 2. Results and Discussion

### 2.1 The rationale in searching for discoveries at low sampling numbers as an alternative approach

To underline some of the pitfalls of employing traditional methods of statistics at sampling numbers, lower than required, we will use a trivial experiment as a first example. Let’s suppose that one tests the bias of a coin by flipping it *5* times and obtaining only heads. The hypothesis that the coin is fair can be rejected at the *α=0.05* significance level 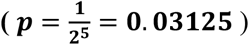. Let’s now assume that m = 10,000 of such experiments are run simultaneously and all coin-flipping results in heads. Applying blindly the Bonferroni method, one concludes that none of the involved coins can be proven to be biased 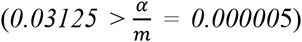. This is a paradoxical result since the probability of obtaining only heads in *50,000* flips of fair coins is abysmally low *(∼10*^*-15,000*^).

Obviously, it can (and it should) be argued that for the Bonferroni correction to work, one should flip each coin at least 18 times (the sampling number provided by Power Analysis). The poor design of the experiment is similar to trying to detect the bias of one coin in a single experiment by flipping it only 3 times. In this case, the level of significance of *0.05* will never be reached.

The conclusion is that the above multiple-comparison experiment was designed poorly and it does not have the required power to detect statistically significant discoveries. However, sometimes the number of required “coin flippings” might simply not be available (e.g., in preliminary data), and the question is what can be inferred from such poorly designed experiments.

Our approach is to, alternatively, look at the likelihood of the overall experiment, assuming that all coins are fair. When *m* uniformly random numbers are generated in the interval [0,1], the probability that exactly *k* of them are less than α is a Poisson distribution, ?*(k, λ)*, with *λ=αm*. Therefore, for the above experiment with fair coins, *m* = 10,000 and *α* = 0.05, one would expect about 500± 22 false discoveries. The probability of the result (all *m* of the *p*-values are less than 0.05), when all of the coins would be fair is∼ ?(10000, 500). It is a truly “black swan” event of the order of *450 σ*. Therefore, whereas no individual hypothesis can be rejected (in the traditional sense) at the significance level *α*, accepting all null hypotheses will lead to the acceptance of an extremely unlikely outcome of the whole experiment.

### 2.2 A case study of the likelihood of discoveries

A more realistic example is the experiment from the seminal paper of Benjamini and Hochberg [4]. Neuhhaus et al. [11] investigated the effects of administrating a new front-loaded recombinant tissue-type plasminogen activator (rt-PA) in comparison with a standard regimen of an isolated plasminogen streptokinase activator (APSAC). They found the effects to be statistically significant to a 0.05 level (*p=*0.0095). However, it was ignored that multiple hypotheses (i.e., 15 hypotheses) were tested simultaneously. When that was considered, the Bonferroni correction rendered the result not statistically significant [4]. Benjamini and Hochberg made justice to the original claim, showing that their correction (controlling only *FDR* at *α*) suggests that the result, however, it actually is statistically significant [4].

Another point worth considering is the likelihood of obtaining 9 *p*-values lower than 0.05 in a 15 comparisons experiment, if only *k* true discoveries are present. This can be determined easily via the Poisson distribution, as described above, and the corresponding probabilities are plotted in Figure 1.

**Figure 1.**
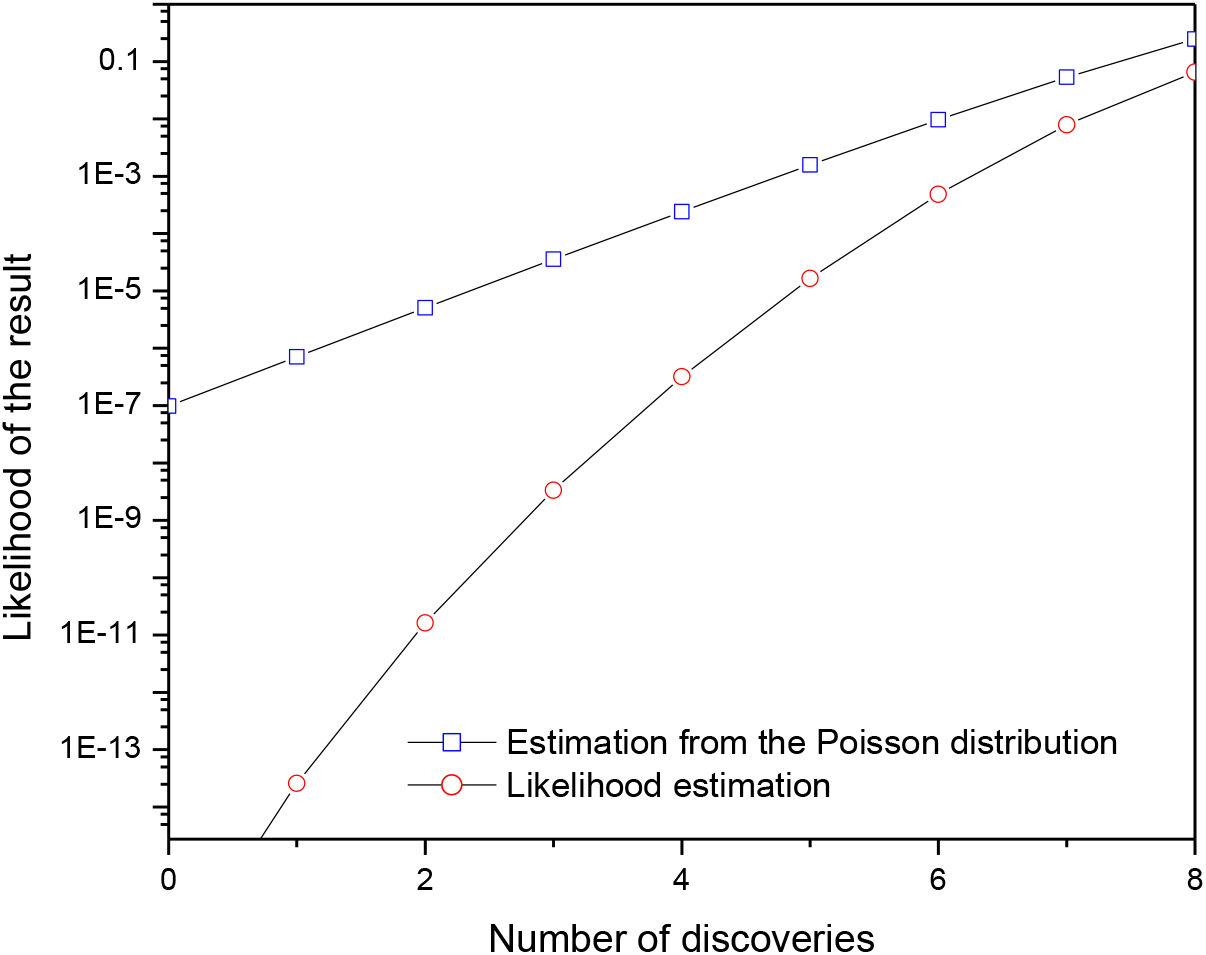
The likelihood of obtaining the result when only k discoveries are present.

This intuitive approach is very simplistic and carries a lot of incertitude. For example, one can alternatively look for the probability of obtaining 4 *p*-values less than 0.001 when testing 15 hypotheses, and so on. The likelihood calculations can be made more accurately by making the rough assumption that if *p*_*j*_ is the probability of a null hypothesis for comparison *j*, then *1-p*_*j*_ is the probability of a non-null hypothesis.

In an experiment, the likelihood that exactly k=0, 1,…, 15 discoveries exist in the family of 15 hypotheses for a given *p*_*j*,_ can be obtained by:

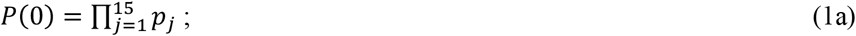

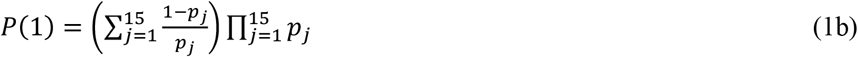

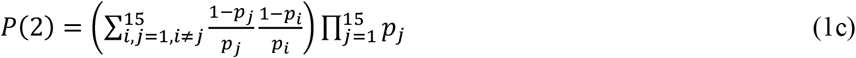

and so on.

These probabilities, which are also plotted in Figure 1, suggest that obtaining such *p*-values, if fewer than 5 discoveries are present, would be unlikely. This neither infirms nor confirms the existence of true discoveries, it just calculates the likelihood that those *p*-values could be obtained as a function of the number of true discoveries existing in the underlying experiment.

Our suggested method for estimating the number of discoveries is based on the assumption that, if no discoveries are present, the obtained *p*-values should be sampled from a uniform random distribution [0,1]. If one discovery exists, the remaining *m-1* of *p*-values should be sampled from a uniform random distribution, and so on. The cumulative function of such a distribution (CDF) is *CDF(x)=x*, and a common test to decide whether the sampled values can be attributed as coming from the underlying distribution is Kologorov-Smirnov [11].

In Figure 2 are plotted the cumulative distribution functions for the obtained *p*-values with the assumption that there are 0 discoveries (a), 3 discoveries (b), and 4 discoveries (c). The Kolmogorov-Smirnov test indicate that the three null hypotheses (that there are only 0, only 3 or only 4 discoveries, respectively), are all rejected at the level of significance of *α*=0.05.

**Figure 2.**
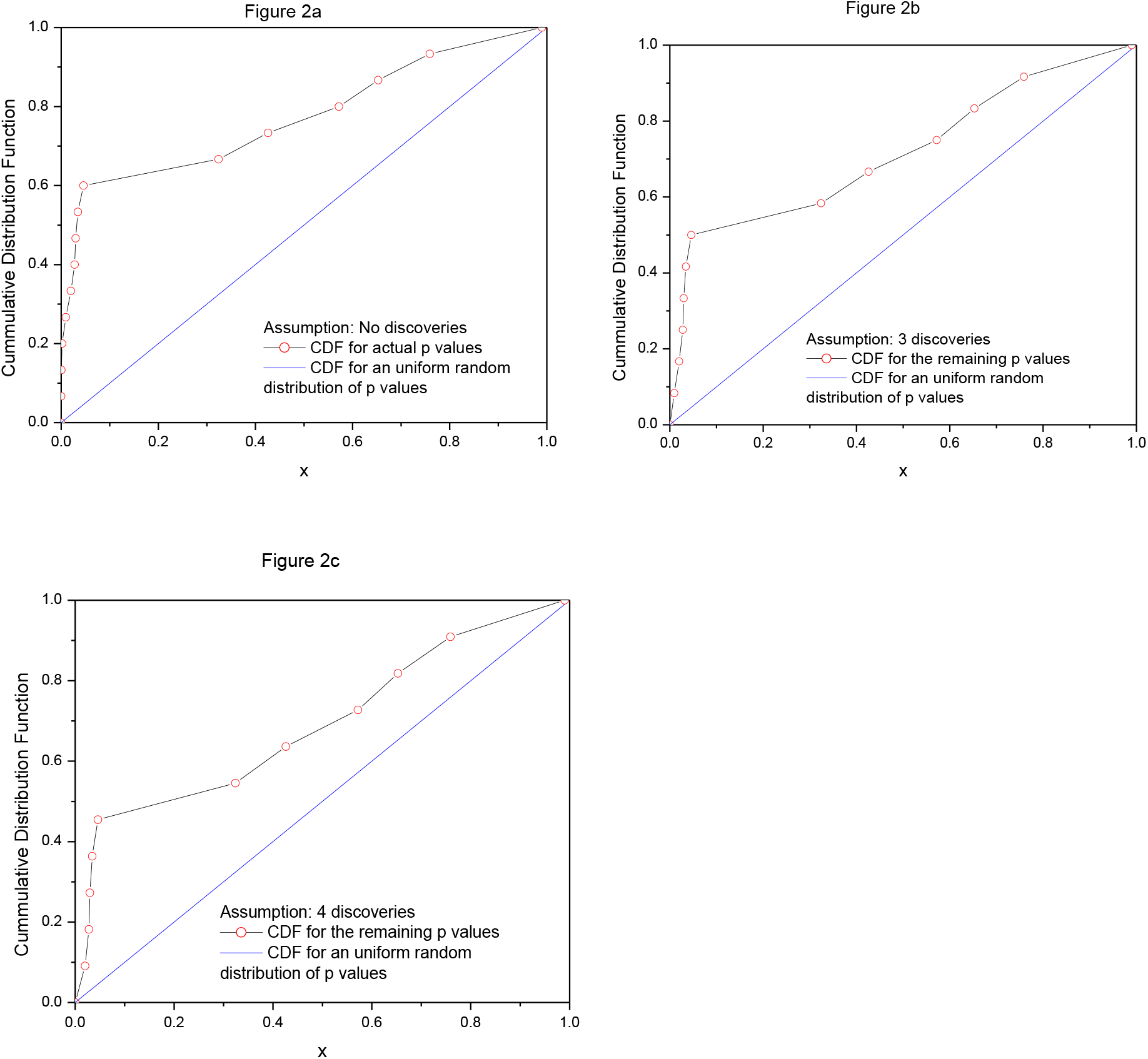
Comparison between the p-values using the CDF of a uniformly random distribution with those from: **(a)** the CDF if no discoveries are present and all of the 15 p-values are uniformly random, **(b)** the CDF of the remaining 12 p-values if 3 discoveries are present, and **(c)** the CDF for the remaining 11 p values if 3 discoveries are present. In all cases, the hypothesis that the p-values are uniformly random is rejected at a 0.05 significance level.

### 2.3 A Monte-Carlo simulation for the number of discoveries in large multiple comparison experiments

We will now test our suggested procedure for evaluating the existing number of discoveries in *1000* simulations of a large number of multi-comparisons (m=10,000) using Monte Carlo. The samplings for the (*m-m*_*0*_) true null hypothesis comparisons and for *m*_*0*_ comparisons are drawn from a standard normal distribution (*μ*_*0*_=0, *σ*=1) and from the true discoveries (*μ*=1, *σ*=1), respectively.

Power analysis implies that if one comparison (*m*=1) is tested, at least 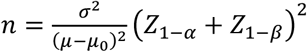 of several samplings should be employed [1]. For *α*=0.05 and β=0.2, the corresponding number is If *m*=10,000 multiple comparisons are tested simultaneously, the replacement of *Z*_1−α_ with 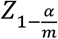 infers that n=38 sampling points are needed. The Z values are calculated from Student’s *t* distribution [1].

The calculations presented here are for *m*_*0*_=1000 discoveries. All of the other values for *m*_*0*_, that were tested by us, led to a similar behavior. At large sampling numbers (*n*=40), as shown in Figure 3a for one of the simulations with *m*=10,000 and *m*_*0*_=1000, the cut-offs for both Bonferroni (B) and Benjamini-Hochberg (BH) methods are very accurate in determining the true number of discoveries in the experiment. As expected, the BH method is more lenient in accepting discoveries than the B method, which overestimates them slightly. A different outcome is seen in Figure 3b for low sampling numbers (*n*=10), where the BH method fails to detect most of the *m*_*0*_ *= 1000* existing discoveries, and the B method fails to detect all of them.

**Figure 3.**
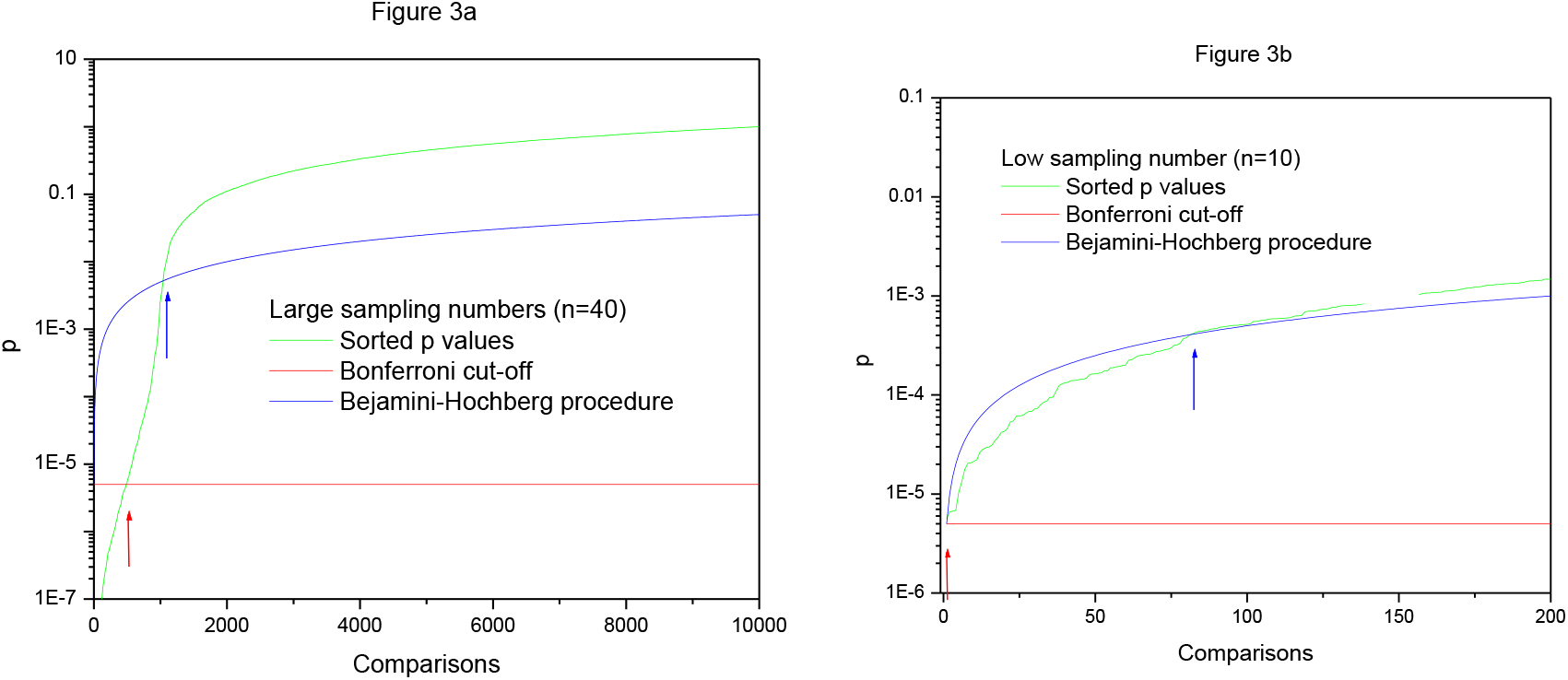
Estimation of the performance of Bonferroni and Benjamini-Hochberg methods: **(a)** at large sampling numbers (n=40) and **(b)** at low sampling numbers (n=10).

A different situation is observed in Figure 4 for the chance that the cumulative function for the *p*-values is similar to the cumulative function for the uniform random distribution, F(x). As shown in this figure, the departure from the cumulative function F(x)=x of a uniform random distribution has a much smaller dependence on the sampling number *n*. For both n=40 (large sampling numbers) and n=10 (low sampling numbers), the departure from the CDF of uniformly random numbers is significant.

**Figure 4.**
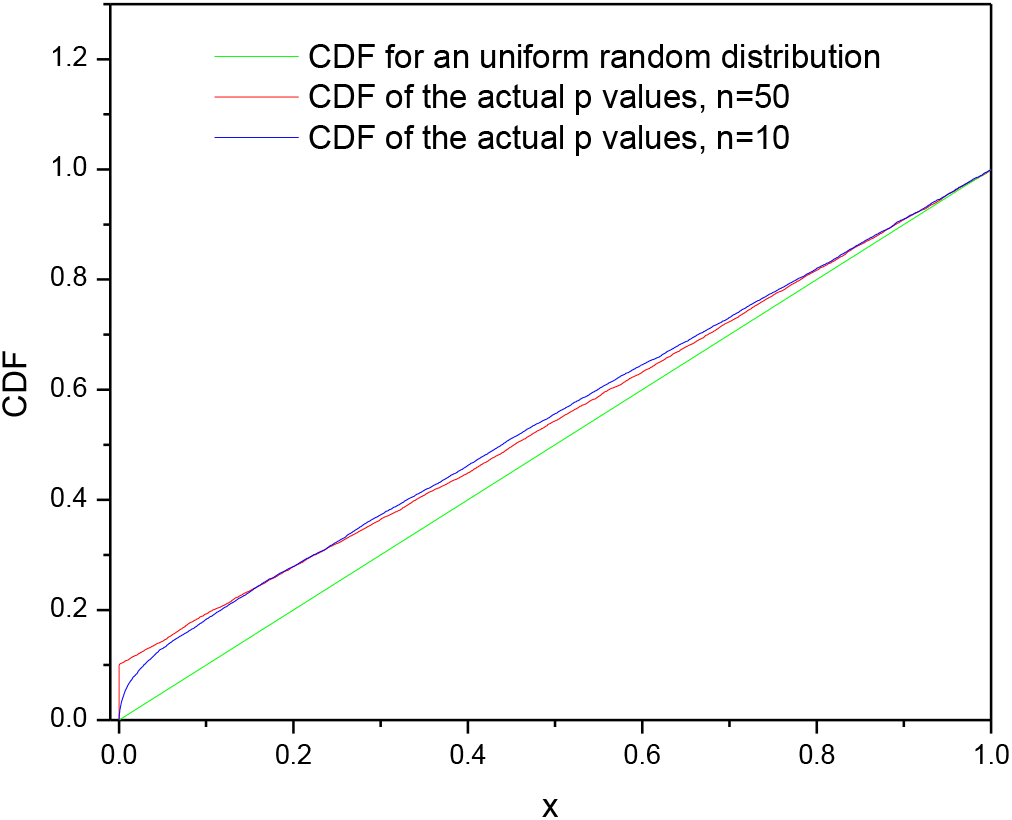
CDF of the p-values showing that the outcome is less affected by the sampling number n.

A combined outcome of the *1000* Monte Carlo simulations is presented in Figure 5 a-f. As observed in Figure 5a, at large sampling numbers (*n*=50), the number of discoveries is well detected by both B and BH methods, with the BH method overestimating them slightly, as expected. At the minimum required sampling number (*n*=40, Figure 5b), the B method detects about 80% of discoveries, while the BH method improves over this estimation rate. However, below the required minimum sampling number, as revealed in Figure 5c for *n*=30 and Figure 5d for *n=*20, the B method fails to identify many existing discoveries. At even lower sampling numbers of *n*=10, the B method fails to identify any discovery, while the BH method fails to identify most of them (Figure 5e). However, the method we proposed here can detect the presence of a majority of discoveries.

**Figure 5.**
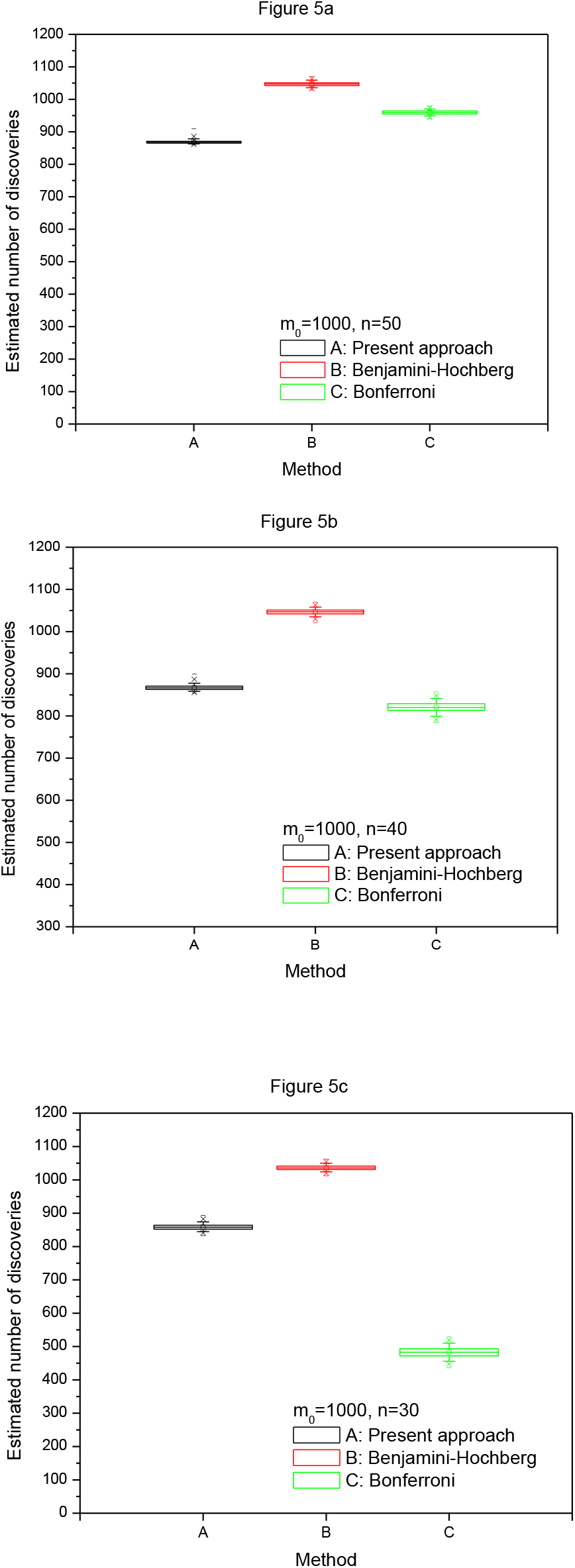
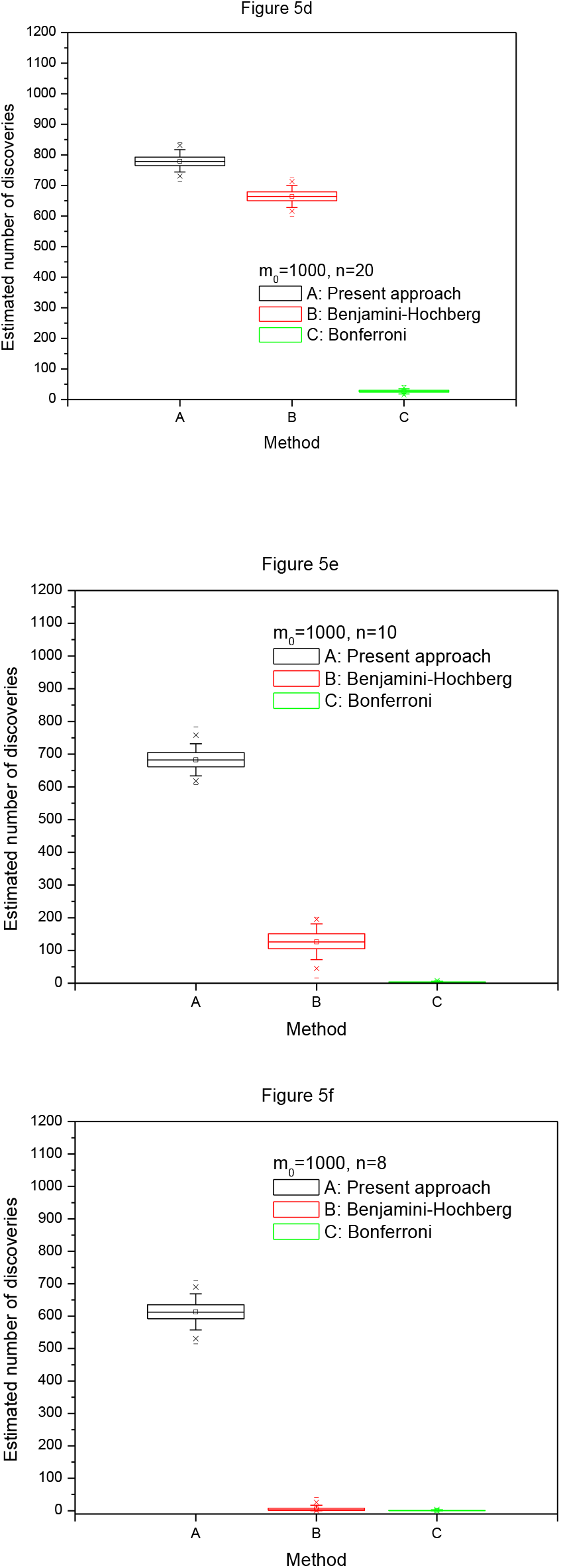
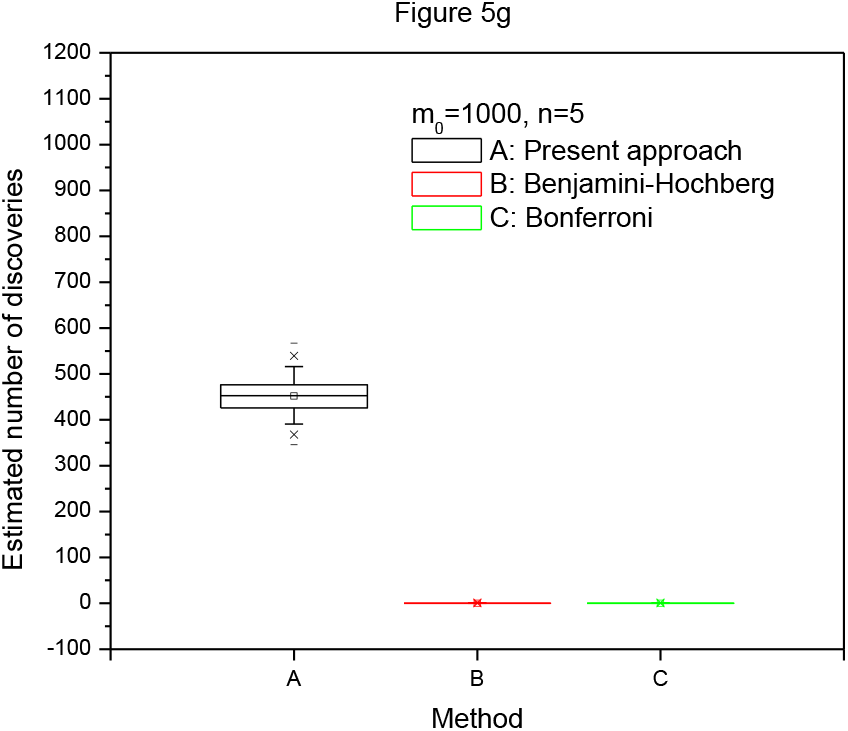
Estimation of the number of discoveries (1000 trials, m=10,000, m_0_=1000), at various sampling numbers (**a:** n=50; **b:** n=40; **c:** n=30; **d:** n=20; **e:** n=10; **f:** n=8; **g:** n=5).

It will be worth determining the ability of our method to correctly identify the existence of discoveries, at very low sample numbers. The numbers employed in Figures 5e and f, *n*=8 and *n*=5, respectively, are too low to be considered in a search for a discovery in a single comparison experiment, let alone in 10,000 multiple comparison experiments. However, the method suggests that a large number of discoveries might exist, while the traditional method fails to identify their presence. This is because the existence of the discoveries affects the uniform random distributions of the *p*-values of the true null hypothesis, even at very low sampling numbers.

## 3. Conclusions

The traditional methods for finding discoveries in multiple comparison experiments can be as accurate as desired if the sampling numbers are sufficiently high. It is well known that they provide a large number of false negatives if the sampling numbers are too low. Increasing the sampling number is time-consuming and costly, and, in some instances cannot be done. An example consists of experiments that have been already completed using low sampling numbers and no options for increasing this number. Furthermore, preliminary multi-comparison experiments, such as qPCR, are performed typically at very low sampling numbers. Thus, it is of interest to estimate the number of likely true discoveries that might exist in such experiments.

The idea behind our proposed method is that in the absence of discoveries, the cumulative function of *p*-values should be similar to the cumulative function of a uniform random distribution. Departure from this distribution can be attributed to the existence of discoveries in the experiment. A Monte Carlo analysis shows that our method is efficient in detecting discoveries even at very low sampling numbers. Therefore, it can be used for secondary analysis of existing data, to indicate the likelihood of discoveries in preliminary data, and for efficient design of further experiments. While the currently employed Bonferroni or Benjamini-Hochberg methods are working well when a required minimum number of sampling is met (their designed framework), there is a need for new methods that allow discoveries at a lower number of samplings where these methods fail. Our simple approach remains powerful for such sampling numbers, for which traditional statistics methods do not provide good results.

## References

[1] Anderson S. Biostatistics: A computing approach. CRC Press; 2011 Dec 20.

[2] Holm S. A simple sequentially rejective multiple test procedure. Scandinavian journal of statistics. 1979 Jan 1:65–70.

[3] Hochberg Y. A sharper Bonferroni procedure for multiple tests of significance. Biometrika. 1988 Dec 1;75(4):800–2.

[4] Benjamini Y, Hochberg Y. Controlling the false discovery rate: a practical and powerful approach to multiple testing. Journal of the royal statistical society. Series B (Methodological). 1995 Jan 1:289–300.

[5] Pike N. Using false discovery rates for multiple comparisons in ecology and evolution. Methods in Ecology and Evolution. 2011 Jun 1;2(3):278–82.

[6] Storey JD. A direct approach to false discovery rates. Journal of the Royal Statistical Society: Series B (Statistical Methodology). 2002 Aug 1;64(3):479–98.

[7] Pounds S, Morris SW. Estimating the occurrence of false positives and false negatives in microarray studies by approximating and partitioning the empirical distribution of p-values. Bioinformatics. 2003 Jul 1;19(10):1236–42.

[8] Pounds S, Cheng C. Robust estimation of the false discovery rate. Bioinformatics. 2006 Jun 15;22(16):1979–87.

[9] L’ecuyer, P. and Simard, R., 2007. TestU01: AC library for empirical testing of random number generators. ACM Transactions on Mathematical Software (TOMS), 33(4), pp.1–40.

[10] Berger, V.W. and Zhou, Y., 2014. Kolmogorov–Smirnov test: Overview. Wiley statsref: Statistics reference online.

[11] Neuhaus, K.L., Von Essen, R., Tebbe, U., Vogt, A., Roth, M., Riess, M., Niederer, W., Forycki, F., Wirtzfeld, A., Maeurer, W. and Limbourg, P., 1992. Improved thrombolysis in acute myocardial infarction with front-loaded administration of alteplase: results of the rt-PA-APSAC patency study (TAPS). Journal of the American College of Cardiology, 19(5), pp.885–891.

